# Interferon Resistance of Emerging SARS-CoV-2 Variants

**DOI:** 10.1101/2021.03.20.436257

**Authors:** Kejun Guo, Bradley S. Barrett, Kaylee L. Mickens, Ezster K. Vladar, James H. Morrison, Kim J. Hasenkrug, Eric M. Poeschla, Mario L. Santiago

## Abstract

The emergence of SARS-CoV-2 variants with enhanced transmissibility, pathogenesis and resistance to vaccines presents urgent challenges for curbing the COVID-19 pandemic. While Spike mutations that enhance virus infectivity or neutralizing antibody evasion may drive the emergence of these novel variants, studies documenting a critical role for interferon responses in the early control of SARS-CoV-2 infection, combined with the presence of viral genes that limit these responses, suggest that interferons may also influence SARS-CoV-2 evolution. Here, we compared the potency of 17 different human interferons against multiple viral lineages sampled during the course of the global outbreak, including ancestral and four major variants of concern. Our data reveal increased interferon resistance in emerging SARS-CoV-2 variants, suggesting that evasion of innate immunity may be a significant, ongoing driving force for SARS-CoV-2 evolution. These findings have implications for the increased lethality of emerging variants and highlight the interferon subtypes that may be most successful in the treatment of early infections.

**Author Summary:** In less than 2 years since its spillover into humans, SARS-CoV-2 has infected over 220 million people, causing over 4.5 million COVID-19 deaths. High infection rates provided substantial opportunities for the virus to evolve, as variants with enhanced transmissibility, pathogenesis, and resistance to vaccine-elicited neutralizing antibodies have emerged. While much focus has centered on the Spike protein which the virus uses to infect target cells, mutations were also found in other viral proteins that might inhibit innate immune responses. Specifically, viruses encounter a potent innate immune response mediated by the interferons, two of which, IFNα2 and IFNβ, are being repurposed for COVID-19 treatment. Here, we compared the potency of human interferons against ancestral and emerging variants of SARS-CoV-2. Our data revealed increased interferon resistance in emerging SARS-CoV-2 strains that included the alpha, beta, gamma and delta variants of concern, suggesting a significant, but underappreciated role for innate immunity in driving the next phase of the COVID-19 pandemic.

## Results

The human genome encodes a diverse array of antiviral interferons (IFNs). These include the type I IFNs (IFN-Is) such as the 12 IFNα subtypes, IFNβ and IFNω that signal through ubiquitous IFNΑR receptor, and the type III IFNs (IFN-IIIs) such as IFNλ1, IFNλ2 and IFNλ3 that signal through the more restricted IFNλR receptor that is present in lung epithelial cells [1]. IFN diversity may be driven by an evolutionary arms-race in which viral pathogens and hosts reciprocally evolve countermeasures [2]. For instance, the IFNα subtypes exhibit >78% amino acid sequence identity, but IFNα14, IFNα8 and IFNα6 most potently inhibited HIV-1 *in vitro* and *in vivo* [3–5], whereas IFNα5 most potently inhibited influenza H3N2 in lung explant cultures [6]. Even though SARS-CoV-2 was sensitive to IFNα2, IFNβ, and IFNλ [7–9], and clinical trials of IFNα2 and IFNβ demonstrated therapeutic promise for COVID-19 [10–12], a direct comparison of multiple IFN-Is and IFN-IIIs against diverse SARS-CoV-2 variants of concern has not yet been done.

The current study was initially undertaken to determine which IFNs would best inhibit SARS-CoV-2. These first set of experiments were performed between December 2020 and March 2021, and we selected 5 isolates from prominent lineages [13] during this phase of the pandemic (Fig 1, S1 Table). USA-WA1/2020 is the standard strain utilized in many *in vitro* and *in vivo* studies of SARS-CoV-2 and belongs to lineage A [13]. It was isolated from the first COVID-19 patient in the US, who had a direct epidemiologic link to Wuhan, China, where the virus was first detected [14]. By contrast, subsequent infection waves from Asia to Europe [15] were associated with the emergence of the D614G mutation [16]. Lineage B strains with G614 spread globally and displaced ancestral viruses with striking speed, likely due to increased transmissibility [17, 18]. These strains accumulated additional mutations in Italy as lineage B.1 which then precipitated a severe outbreak in New York City [19]. Later in the United Kingdom (U.K.), lineage B.1.1.7 acquired an N501Y mutation associated with enhanced transmissibility [13]. Lineage B.1.351, first reported in South Africa, additionally acquired an additional E484K mutation associated with resistance to neutralizing antibodies [20, 21]. Both B.1.1.7 and B.1.351 were reported in multiple countries and in some cases have become dominant for extended periods [22]. We obtained representative SARS-CoV-2 isolates of the B, B.1, B.1.1.7 and B.1.351 lineages (S1 Table). Each stock was sourced from beiresources.org and amplified once in a human alveolar type II epithelial cell line (A549) that we have stably transduced with the receptor ACE2 (A549-ACE2) (S1A Fig).

**Figure 1.**
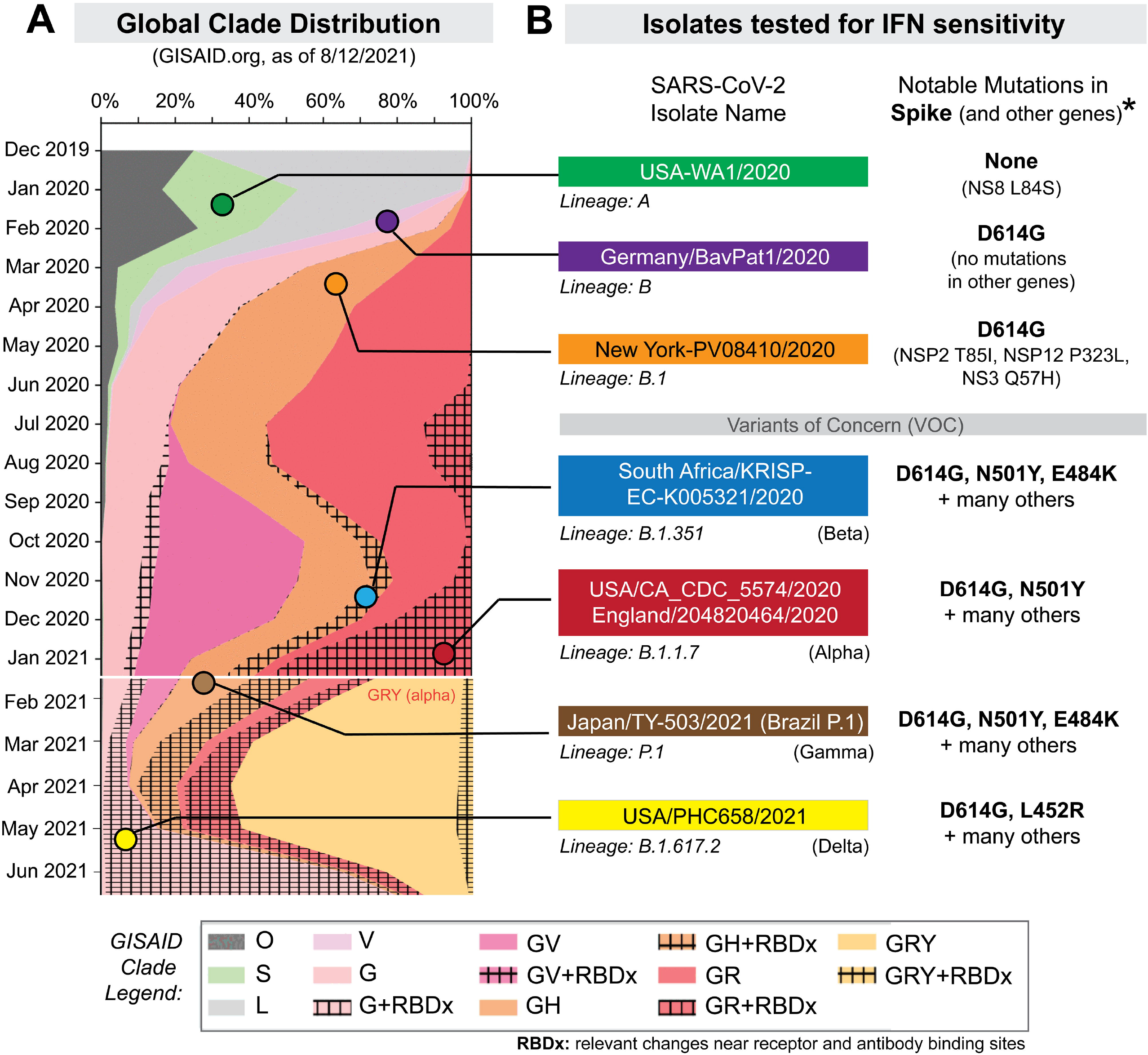
Selection of SARS-CoV-2 strains for IFN sensitivity studies. (A) Global distribution of SARS-CoV-2 clades. GISAID.org plotted the proportion of deposited sequences in designated clades against collection dates. The six isolates chosen are noted by colored dots. (B) SARS-CoV-2 strains selected for this study included representatives of lineages A, B, B.1, B.1.351 and B.1.1.7 (S1 Table). Lineage P.1 (which branched off from lineage B.1.1.28) and B.1.617.2 were added after the initial manuscript submission; and was evaluated for IFNβ and IFNλ1 sensitivity. Lineage B isolates encode the D614G mutation associated with increased transmissibility. Note that the B.1.1.7 strain was later updated to belong to the GISAID clade, ‘GRY’. *Amino acid mutations were relative to the reference hCOV-19/Wuhan/WIV04/2019 sequence.

A549-ACE2 cells were pre-incubated with 17 recombinant IFNs (PBL Assay Science) overnight in parallel and in triplicate, then infected with a non-saturating virus dose for 2 h (S1B Fig). We normalized the IFNs based on molar concentrations similarly to our previous work with HIV-1 [3, 23]. To enable high-throughput evaluation of the antiviral activities of the numerous IFNs against the multiple live SARS-CoV-2 isolates, we used a quantitative PCR (qPCR) assay to determine amounts of virus produced 24 hours after infection (Fig 2A). Initial dose-titrations showed that a 2 pM concentration fell within the dynamic range of activity and maximally distinguished the antiviral activities of IFNs with widely divergent potencies, i.e., IFNβ and IFNλ1 (S1C Fig). Of note, the IFNβ and IFNλ1 doses used did not significantly affect cell viability (S1D Fig). Thus, 2 pM doses were used for additional antiviral activity testing. We also evaluated the qPCR assay against a VeroE6 plaque assay using triplicate serial dilutions of a SARS-CoV-2 isolate (B.1.351). Virus titers obtained using these two assays were strongly correlated (S2A Fig). However, the VeroE6 plaque assay had ~2-log lower dynamic range; we estimate that 1 plaque forming unit corresponds to ~900 SARS-CoV-2 N1 copies (S2A Fig). Virus copy numbers also correlated with the numbers of primary airway epithelial cells infected with different SARS-CoV-2 variants as quantified by immunofluorescence (S2B Fig). Thus, we employed the qPCR assay to robustly distinguish the antiviral activity of the different interferons.

**Figure 2.**
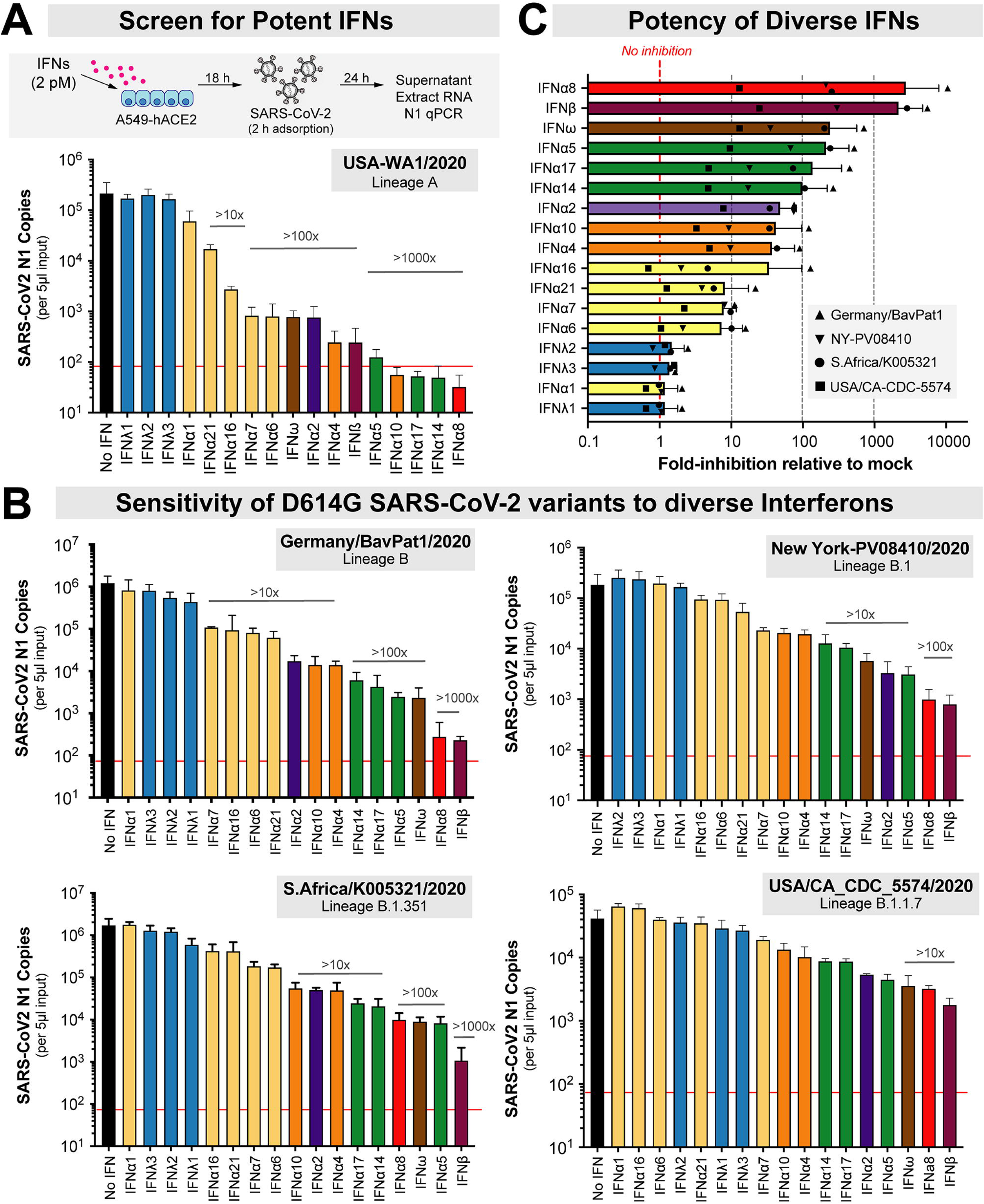
Sensitivity of SARS-CoV-2 strains to IFN-I and IFN-III interferons. (A) Antiviral assay using recombinant IFNs (2 pM) in A549-ACE2 cells. The red line corresponds to the qPCR detection limit (90 copies/reaction, or 1.8 x 10^4^ copies/ml). (B) Viral copy numbers in D614G+ isolates, showing a similar rank-order of IFNs from least to most potent. (C) The average fold-inhibition relative to mock for lineage B, B.1, B.1.351 and B.1.1.7 isolates are shown. The most potent IFNs are shown top to bottom. For all panels, bars and error bars correspond to means and standard deviations.

In the absence of IFN, all 5 isolates reached titers of ~10^4^-10^6^ copies per 5 μl input of RNA extract (Fig 2). Using absolute copy numbers (Fig 2) or values normalized to mock as 100% (S2 Fig), the 17 IFNs showed a range of antiviral activities against SARS-CoV-2. The 3 IFNλ subtypes exhibited none to very weak (<2-fold) antiviral activities compared to most IFN-Is (Fig 2 and S3 Fig, blue bars). This was despite the fact that the assay showed a robust dynamic range, with some IFNs inhibiting USA-WA1/2020 >2500-fold to below detectable levels (Fig 2A). IFN potencies against the 5 isolates correlated with each other (S4 Fig), and a similar rank-order of IFN antiviral potency was observed for G614+ isolates (Fig 2B, S3 Fig). Overall, IFNα8, IFNβ and IFNω were the most potent, followed by IFNα5, IFNα17 and IFNα14 (Fig 2C); the type III (λ) IFNs were least potent.

The molecular basis for the diverse antiviral effects of the highly related IFNα subtypes has been an active area of investigation, particularly with regard to the relative contributions of quantitative (signaling) versus qualitative (differential gene regulation) mechanisms [2–5]. We reported that inhibition of HIV-1 by the IFNα subtypes correlated with IFNΑR signaling capacity and binding affinity to the IFNΑR2 subunit [3, 23]. IFNΑR signaling capacity, as measured in an IFN-sensitive reporter cell line (iLite cells; Euro Diagnostics), correlated with the antiviral potencies of the IFNα subtypes against SARS-CoV-2 lineages A and B, but not B.1, B.1.351 or B.1.1.7 (Fig 3A). IFNAR binding affinities as measured by surface plasmon resonance by the Schreiber group [24] did not correlate with IFNα subtype inhibition of SARS-CoV-2 (Fig 3B). As the recombinant IFNs used in this study was from the same source as that of the prior HIV-1 study [3, 23], we also determined if the IFNs that potently inhibit HIV-1 also function similarly against SARS-CoV-2. Notably, the correlations between SARS-CoV-2 and HIV-1 inhibition [3] were weak at best (Fig 3C). These findings suggested that IFN-mediated control of SARS-CoV-2 isolates may be qualitatively distinct from that of HIV-1.

**Figure 3.**
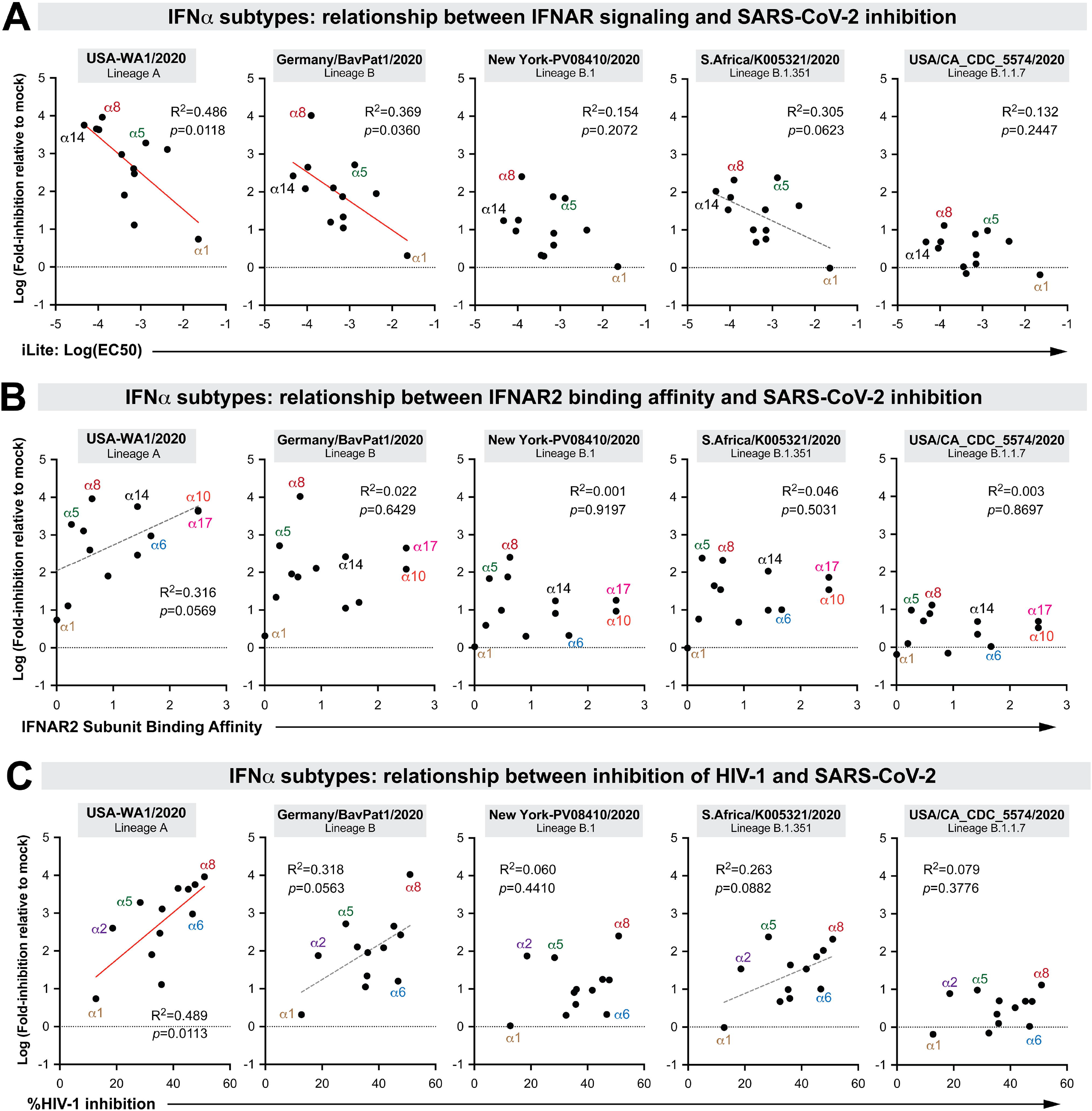
Correlation between SARS-CoV-2 inhibition and biological properties of IFN α subtypes. Log-transformed IFN-inhibition values relative to mock for the 5 different SARS-CoV-2 strains were compared to previously published values on (A) 50% effective concentrations in the iLite assay, a reporter cell line encoding the IFN sensitive response element of *ISG15* linked to firefly luciferase [23]; (B) IFNΑR2 subunit binding affinity, as measured by surface plasmon resonance by the Schreiber group [24]; and (C) HIV-1 inhibition values, based on % inhibition of HIV-1 p24+ gut lymphocytes relative to mock as measured by flow cytometry [3]. Each dot corresponds to an IFNα subtype. Linear regression was performed using GraphPad Prism 8. Significant correlations (*p*<0.05) were highlighted with a red best-fit line; those that were trending (*p*<0.1) had a gray, dotted best-fit line.

We generated a heat-map to visualize the antiviral potency of diverse IFNs against the 5 isolates and observed marked differences in IFN sensitivities (Fig 4A). Pairwise analysis of antiviral potencies between isolates collected early (January 2020) and later (March-December 2020) during the pandemic were performed against the 14 IFN-Is (IFN-III data were not included due to low antiviral activity, Fig. 2). The overall IFN-I sensitivity of USA-WA1/2020 and Germany/BavPat1/2020 isolates were not significantly different from each other (Fig 4B). In contrast, relative to Germany/BavPat1/2020, we observed 17 to 122-fold IFN-I resistance of the emerging SARS-CoV-2 variants (Fig 4C), with the B.1.1.7 strain exhibiting the highest IFN-I resistance (this can also be seen in Fig. 3). The level of interferon resistance was especially striking when compared to the early pandemic USA-WA1/2020 strain, where emerging SARS-CoV-2 variants exhibited 25 to 322-fold higher IFN-I resistance (Fig 4D).

**Figure 4.**
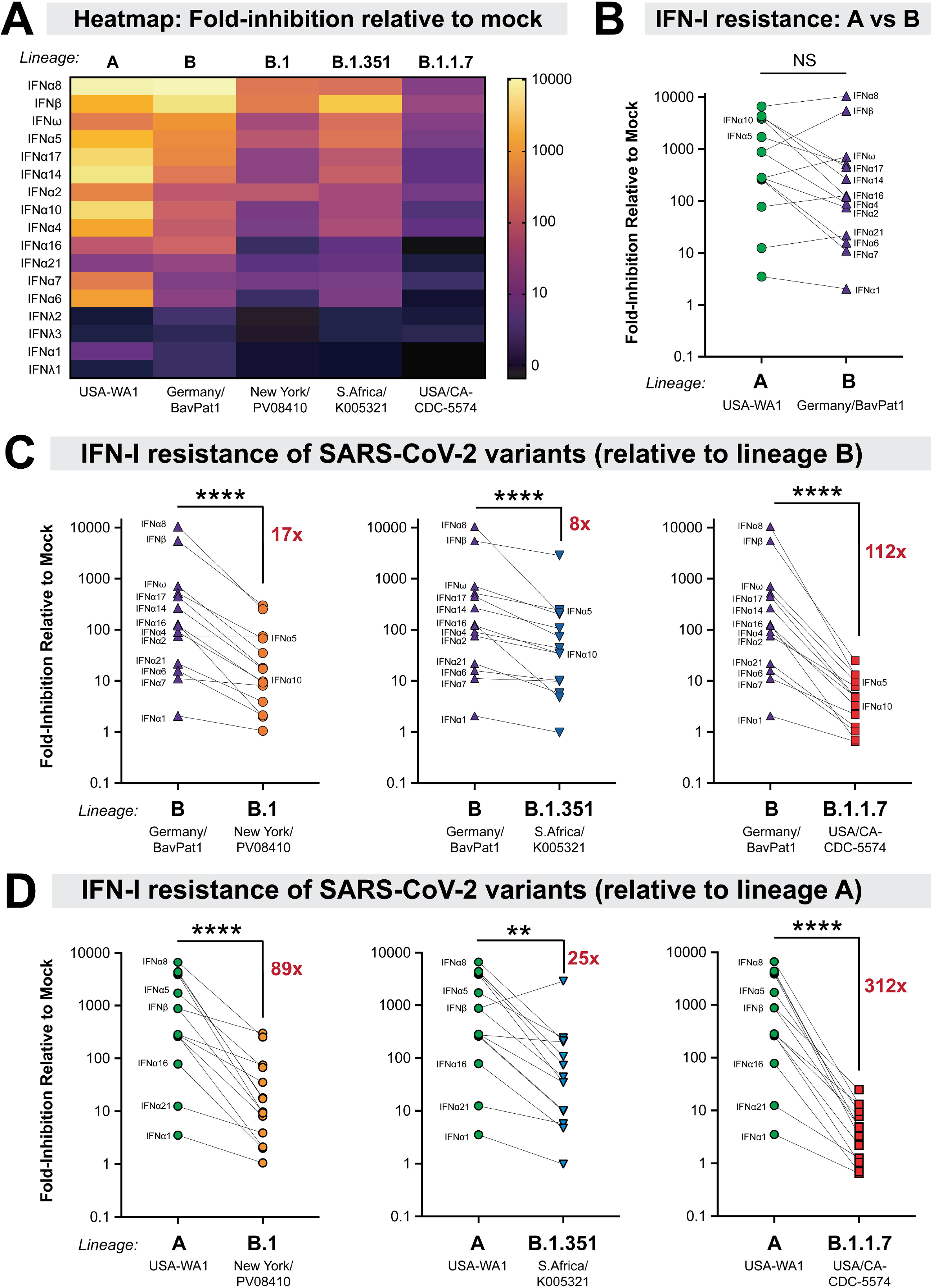
Increased IFN-I resistance of emerging SARS-CoV-2 variants. (A) Heatmap of fold-inhibition of representative strains from the lineages noted. Colors were graded on a log-scale from highest inhibition (yellow) to no inhibition (black). Comparison of IFN-I sensitivities between (B) lineage A and B isolates; (C) lineage B versus B.1, B.1.351 and B.1.1.7 and (D) lineage A versus B.1, B.1.351 and B.1.1.7. The mean fold-inhibition values relative to mock were compared in a pairwise fashion for the 14 IFN-Is. In (C) and (D), the average fold-inhibition values were noted. Differences were evaluated using a nonparametric, two-tailed Wilcoxon matched-pairs signed rank test. NS, not significant; ****, *p*<0.0001.

The experiments to this point allowed for the simultaneous analysis of 17 IFNs against multiple SARS-CoV-2 isolates, but did not provide information on how different IFN-I doses affect virus replication. It also remained unclear if the emerging variants were resistant to IFN-IIIs. We therefore titrated a potent (IFNβ; 0.002 to 200 pM) and a weak (IFNλ1; 0.02 to 2000 pM) interferon against the lineage A, B, B.1, B.1.1.7 and B.1.351 viruses (Fig 5 and S5 Fig). Of note, as the pandemic progressed in the past year, new variants of concern (VOCs) became dominant in several countries; the WHO implemented a simplified Greek letter nomenclature for these VOCs. We therefore included 3 additional VOCs, which were also obtained from the BEI repository: (1) a second B.1.1.7 (alpha) isolate, England/204820464/2020; (2) an isolate from lineage P.1 (gamma), which branched off from lineage B.1.1.28; and (3) an isolate from lineage B.1.617.2 (delta) (S1 Table). Lineage P.1 was first described in an outbreak of SARS-CoV-2 in Manaus, Brazil, which occurred in a population with high levels of prior infection. P.1 independently acquired the E484K mutation [25, 26] (Fig 1A, S1 Table). The delta strain was first reported in India in early 2021 [27, 28], and as of July 2021, has become the dominant variant worldwide, including the USA [29]. The delta strain was particularly concerning as it was frequently observed in breakthrough infections among fully-vaccinated individuals [30, 31].

**Figure 5.**
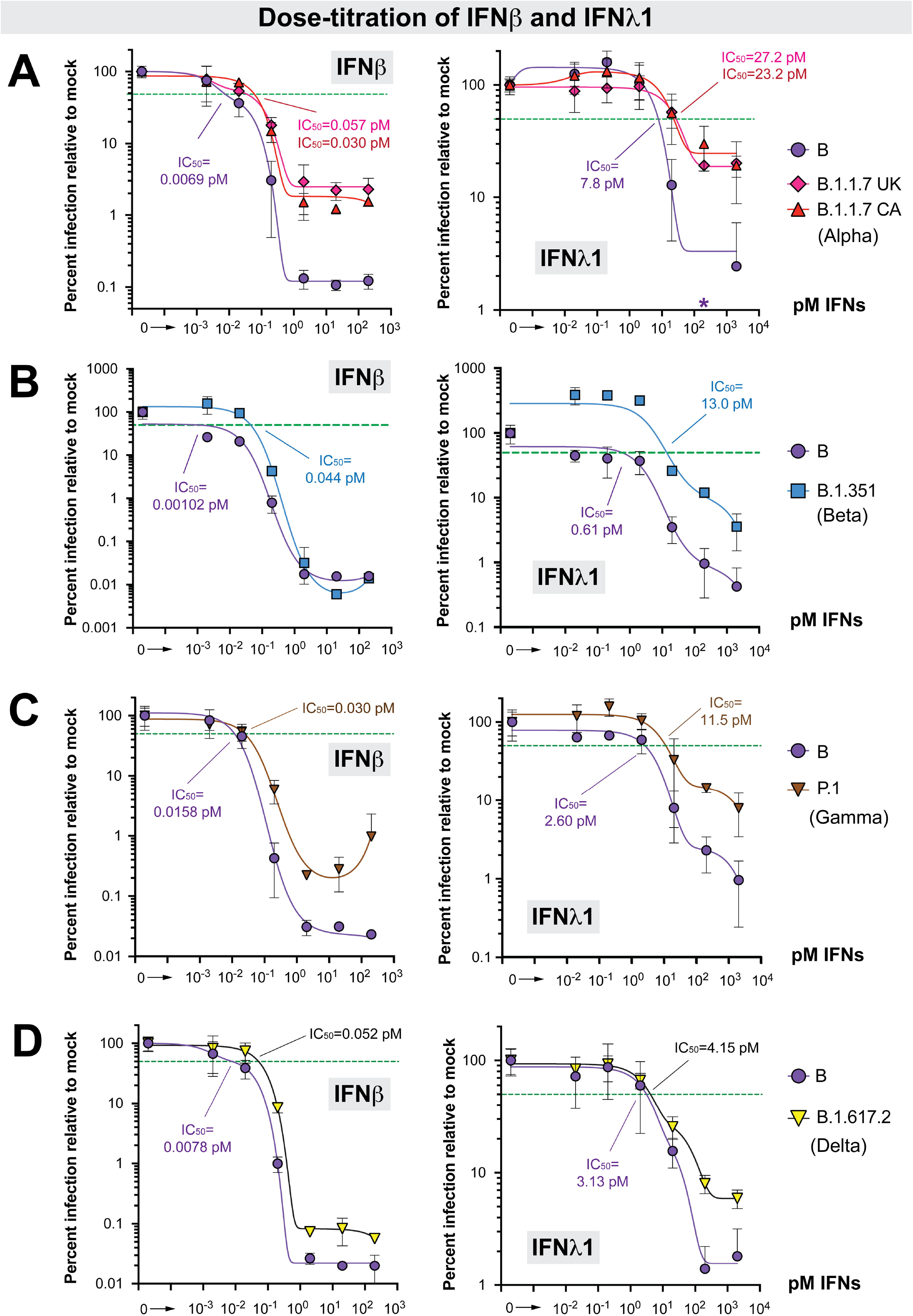
Dose-titration of ancestral lineage B versus four variants of concern against IFN β and IFN λ1. Data from four separate experiments (panels A-D) are shown. (A) Dose-titration of IFNβ and IFNλ1 against lineage B (Germany/BavPat1/2020) versus B.1.1.7 (alpha) isolates. In addition to USA/CA_CDC_5574/2020, we also evaluated a second B.1.1.7 isolate from the United Kingdom (UK), England/204820464/2020. *The value at 200 pM IFNλ1 for the lineage B isolate was 0.54, precluding efforts for finding a best-fit curve for IC50 determination; this datapoint was therefore not included in the curve fitting. (B) IC50 comparison between a lineage B (Germany/BavPat1/2020) and a B.1.351 (beta) isolate (South Africa/KRISP-EC-K005321/2020). (C) IC50 comparison between a lineage B isolate (Germany/BavPat1/2020) and a P.1 (gamma) isolate (Japan/TY7-503/2021). (D) IC50 comparison between a lineage B isolate (Germany/BavPat1/2020) and a B.1.617.2 (delta) isolate (USA/PHC658/2021). For all panels, A549-ACE2 cells were pre-treated with serial 10-fold dilutions of IFNs for 18 h in triplicate and then infected with SARS-CoV-2. Supernatants were collected after 24 h, SARS-CoV-2 N1 copy numbers were determined by qPCR in triplicate, and then the mean copy numbers were normalized against mock as 100%. Error bars correspond to standard deviations. Non-linear best-fit regression curves of mean normalized infection levels were used to interpolate 50% inhibitory concentrations (green dotted lines).

The lineage A and B isolates were similarly inhibited by IFNβ and IFNλ1 (S5A Fig). Comparing B to B.1, the 50% inhibitory concentration (IC_50_) of the B.1 isolate was 2.6 and 5.5-fold higher IC_50_ for IFNλ1 and IFNβ, respectively (S5B Fig). Comparing B to B.1.1.7, the B.1.1.7 variants IC_50_s were 4.3 to 8.3-fold higher for IFNβ and 3.0 to 3.5 higher for IFNλ1 (Fig 5A). Interestingly, maximum inhibition was not achieved with either IFNβ or IFNλ1 against the B.1.1.7 variant, plateauing at 15 to 20-fold higher levels than the ancestral lineage B isolate (Fig. 5A), which was in sharp contrast to the lineage B.1 isolate (S5B Fig). In a separate experiment, the B.1.351 variant was also more resistant to IFNβ (43-fold) and IFNλ1 (26-fold) compared to the lineage B isolate (Fig 5B). Here, however, maximum inhibition was achieved with IFNβ. The P.1 variant also exhibited higher resistance to IFNβ (1.9-fold) and IFNλ1 (4.4-fold), and the plateau concentration for antiviral activity was >10-fold higher for IFNβ than for the lineage B isolate (Fig. 5C). Consistent with the findings with the other VOCs, the B.1.617.2 (delta) variant was also more resistant to IFNβ (6.7-fold) (Fig. 5D). Although similar IC50s were obtained with IFNλ1, the B.1.617.2 isolate had higher residual replication at the highest doses than the ancestral lineage B isolate (Fig. 5D).

Two months after our initial preprint [32], Thorne *et* al posted data that in Calu-3 cells, a B.1.1.7 isolate, was more resistant to IFNβ than a ‘first wave’ lineage B isolate [33]. We found that lineage A and B isolates replicated poorly in Calu-3 cells, making these cells unsuitable for IFN resistance comparisons between ancestral versus emerging variants (S6A Fig). This was in sharp contrast to A549-ACE2 cells, where we observed high levels of virus production (>10^5^ copies) of all strains studied (S1B Fig). Notably, comparable titers were obtained between the B.1 and B.1.1.7 isolates in Calu-3 cells (S6A Fig). In these cells, the B.1.1.7 isolate was 50-fold more resistant to IFNλ1 than the B.1 isolate (S6B Fig). We also demonstrate that the B.1.1.7 and B.1.617.2 isolates were more resistant to IFNβ than the B.1 isolate (S6C Fig). Altogether, our data demonstrate that the B.1, B.1.1.7, B.1.351, P.1 and B.1.617.2 isolates have evolved to resist the IFN-I and IFN-III response.

## Discussion

Numerous studies have shown that interferons are important for host defense against SARS-CoV-2. This sarbecovirus is believed to have recently crossed the species barrier to humans, either directly from bats or via an intermediate mammalian host(s) [34]. Here, we demonstrate that SARS-CoV-2 has in fact evolved after host switching to become more resistant to human interferons. Moreover, we establish an order of antiviral potency for the diverse type I and III IFNs. IFNλ initially showed promise as an antiviral that can reduce inflammation [35], but our data suggest that for SARS-CoV-2, higher doses of IFNλ may be needed to achieve a similar antiviral effect *in vivo* as the IFN-Is. Nebulized IFNβ showed potential as a therapeutic against COVID-19 [11], and our data confirm IFNβ is highly potent against SARS-CoV-2. However, IFNβ was also linked to pathogenic outcomes in chronic mucosal HIV-1 [23], murine LCMV [36] and if administered late in mice, SARS-CoV-1 and MERS-CoV [37, 38] infection. We previously reported that IFNβ upregulated 2.4-fold more genes than individual IFNα subtypes, suggesting that IFNβ may induce more pleiotropic effects [23]. Among the IFNα subtypes, IFNα8 showed similar anti-SARS-CoV-2 potency as IFNβ. IFNα8 also exhibited high antiviral activity against HIV-1 [3], raising its potential for treatment against both pandemic viruses. Notably, IFNα8 appeared to be an outlier in this regard, as the antiviral potencies of the IFNα subtypes against SARS-CoV-2 and HIV-1 generally did not strongly correlate (Fig. 3C). IFNα6 potently restricted HIV-1 [3, 4] but was one of the weakest IFNα subtypes against SARS-CoV-2. Conversely, IFNα5 strongly inhibited SARS-CoV-2, but weakly inhibited HIV-1 [3]. This lack of correlation is a key point for future studies. Of note, the high potency of IFNα5 and low potency of IFNα6 against an isolate of SARS-CoV-2 (not a variant of concern) were corroborated by another group [39]. Collectively, these data strengthen the theory that diverse IFNs may have evolved to restrict distinct virus families [2, 23]. The mechanisms underlying these interesting qualitative differences remain unclear. While IFNΑR signaling contributes to antiviral potency [3, 4, 24], diverse IFNs may have distinct abilities to mobilize antiviral effectors in specific cell types. Comparing the interferomes induced by distinct IFNs in lung epithelial cells [39] may be useful in prioritizing further studies on this point.

Most significantly, our data reveal for the first time the concerning trend for SARS-CoV-2 variants emerging later in the pandemic – in the setting of prolific replication of the virus in human populations – to resist the antiviral interferon response. Prior to the present work, the emergence and fixation of variants was linked to enhanced viral infectivity and/or neutralizing antibody evasion due to mutations in the Spike protein [13, 16-18, 40]. However, previous studies with HIV-1 suggested that interferons also can shape the evolution of pandemic viruses [41, 42]. In fact, SARS-CoV-2 infected individuals with either genetic defects in IFN signaling [43] or IFN-reactive autoantibodies [44] had increased risk of developing severe COVID-19. As interferons are critical in controlling early virus infection levels, IFN-resistant SARS-CoV-2 variants may produce higher viral loads that could in turn promote transmission and/or exacerbate pathogenesis. Consistent with this hypothesis, some reports have linked B.1.1.7 with increased viral loads [45, 46] and risk of death [47–49]. Notably, infection with B.1.617.2 may yield even higher viral loads than that B.1.1.7 [50].

In addition to Spike, emerging variants exhibit mutations in nucleocapsid, membrane and nonstructural proteins NSP3, NSP6 and NSP12 (S1 Table). In the case of some early pandemic viruses that pre-dated the emergence of the variants of concern, these viral proteins were reported to antagonize IFN signaling in cells [51–53]. To specifically map the virus mutations driving IFN-I resistance in emerging variants, it will be important to generate recombinant viruses to isolate specific mutations, singly or in combination, and individually test candidate single viral protein antagonists as well. This would help to confirm, for example, that the D3L mutation in the B.1.1.7 nucleocapsid may facilitate innate immune evasion by increasing the expression of an interferon antagonist, ORF9b [33]. The nucleocapsid D3L mutation was not observed in the B.1.351, P.1 and B.1.617.2 lineages (S1 Table), which exhibited IFN-I and IFN-III resistance in our experiments. B.1.617.2 (delta) has now replaced B.1.1.7 (alpha) as the dominant strain in many countries [27, 29], but delta did not seem to be any more interferon-resistant than alpha in both A549-ACE2 and Calu-3 cells. Notably, the delta isolate we studied here had a deletion in ORF7a, which may counteract interferon signaling [52]; this deletion was not a cell culture artifact as it was also observed in the clinical isolate. Analysis of delta isolates with or without the ORF7a deletion would be needed to determine whether innate immune evasion may be a factor for why the delta VOC has overtaken other lineages. Future studies should facilitate understanding the molecular mechanisms of interferon resistance, its consequences for COVID-19 pathogenesis, and the development of novel therapies that augment innate immune defenses against SARS-CoV-2.

Overall, the current study suggested a role for the innate immune response in driving the evolution of SARS-CoV-2 that could have practical implications for interferon-based therapies. Our findings reinforce the importance of continued full-genome surveillance of SARS-CoV-2, and assessments of emerging variants not only for resistance to vaccine-elicited neutralizing antibodies, but also for evasion of the host interferon response.

## Materials and Methods

### Cell lines

A549 cells were obtained from the American Type Culture Collection (ATCC) and cultured in complete media containing F-12 Ham’s media (Corning), 10% fetal bovine serum (Atlanta Biologicals), 1% penicillin/streptomycin/glutamine (Corning). Calu-3 cells were also obtained from ATCC and cultured in DMEM supplemented with 10% fetal bovine serum and 1% penicillin/streptomycin/glutamine (Corning). Both cell lines were maintained at 37°C 5% CO_2_. A549 cells were transduced with codon-optimized human ACE2 (Genscript) cloned into pBABE-puro [54] (Addgene). To generate the A549-ACE2 stable cell line, 10^7^ HEK293T (ATCC) cells in T-175 flasks were transiently co-transfected with 60 μg mixture of pBABE-puro-ACE2, pUMVC, and pCMV-VSV-G at a 10:9:1 ratio using a calcium phosphate method [55]. Forty-eight hours post transfection, the supernatant was collected, centrifuged at 1000×*g* for 5 min and passed through a 0.45 μm syringe filter to remove cell debris. The filtered virus was mixed with fresh media (30% vol/vol) that included polybrene (Sigma) at a 6 μg/ml final concentration. The virus mixture was added into 6-well plates with 5×10^5^ A549 cells/well and media was changed once more after 12 h. Transduced cells were selected in 0.5 μg/ml puromycin for 72 h, and ACE2 expression was confirmed by flow cytometry, western blot and susceptibility to HIV-1ΔEnv/SARS-CoV-2 Spike pseudovirions.

### Virus isolates

All experiments with live SARS-CoV-2 were performed in a Biosafety Level-3 (BSL3) facility with powered air-purifying respirators at the University of Colorado Anschutz Medical Campus. The SARS-CoV-2 stocks were obtained from BEI Resources (www.beiresources.org). S1 Table provides detailed information on the source of the material, the catalogue and lot numbers and virus sequence information of both the clinical and cultured stocks. The viruses were propagated in human A549-ACE2 cells unless indicated and harvested by 72 h to minimize mutations that can occur during passage in cell culture, which were documented particularly in nonhuman primate (Vero) or non-alveolar type II (293T) cell lines [56]. The virus stocks had comparable titers >10^6^ TCID_50_/ml (S1A Fig) except for the two B.1.1.7 strains (CA_CDC_5574/2020 and England/204820464/2020). The contents of the entire vial (~0.5 ml) were inoculated into 3 T-75 flasks containing 3×10^6^ A549-ACE2 cells, except for B.1.1.7 which was inoculated into 1 T-75 flask. The supernatants were collected and spun at 2700×*g* for 5 min to remove cell debris, and frozen at −80°C. The A549-amplified stocks were titered according to the proposed assay format (S1B Fig, Fig 2A). Briefly, 2.5×10^4^ A549-ACE2 cells were plated per well in a 48-well plate overnight. The next day, the cells were infected with 300, 30, 3, 0.3, 0.03 and 0.003 μl (serial 10-fold dilution) of amplified virus stock in 300 μl final volume of media for 2 h. The virus was washed twice with PBS, and 500 μl of complete media with the corresponding IFN concentrations were added. After 24 h, supernatants were collected, and cell debris was removed by centrifugation at 3200×*g* for 5 min.

### Cell viability

To evaluate if the IFN doses affected cell viability, we utilized an MTT assay. 1.5×10^4^ A549-ACE2 cells were plated per well in a 96-well plate and treated with 2000 pM IFNλ1, 2 pM IFNλ1, 200 pM IFNβ, 2 pM IFNβ or untreated. Eight replicates were used per treatment group. As a positive control for cell death, the same number of cells were treated with 30% DMSO. 36 hours after treatment, cell proliferation was assessed using the Vybrant MTT Cell Proliferation Assay Kit (Invitrogen). Media was completely removed from cells and replaced with 100 μl of fresh growth media. 10 μl of 12 mM MTT stock solution was added per well and cells were incubated at 37°C for 4 h. 100 μl SDS-HCl solution was added to each well and mixed thoroughly. After an additional 3 h incubation at 37°C, the absorbance was measured at 570 nm and blank corrected to a media only control.

### SARS-CoV-2 quantitative PCR

For rapid and robust assessments of viral replication, we utilized a real-time quantitative PCR (qPCR) approach. This assay would require less handling of infectious, potentially high-titer SARS-CoV-2 in the BSL3 compared to a VeroE6 plaque assay, as the supernatants can be directly placed in lysis buffer containing guanidinium thiocyanate that would inactivate the virus by at least 4-5 log10 [57]. Importantly, residual IFNs in the culture supernatant could further inhibit virus infection in the VeroE6 plaque assay, compromising the infectious titer read-outs. To measure SARS-CoV-2 levels, total RNA was extracted from 100 μl of culture supernatant using the E.Z.N.A Total RNA Kit I (Omega Bio-Tek) and eluted in 50 μl of RNAse-free water. 5 μl of this extract was used for qPCR. Official CDC SARS-CoV-2 N1 gene primers and TaqMan probe set were used [58] with the Luna Universal Probe One-Step RT-qPCR Kit (New England Biolabs):

Forward primer: GACCCCAAAATCAGCGAAAT
Reverse primer: TCTGGTTACTGCCAGTTGAATCTG
TaqMan probe: FAM-ACCCCGCATTACGTTTGGTGGACC-TAMRA

The sequence of the primers and probes were conserved against the 7 SARS-CoV-2 lineages that were investigated. The real-time qPCR reaction was run on a Bio-Rad CFX96 real-time thermocycler under the following conditions: 55°C 10 mins for reverse transcription, then 95°C 1 min followed by 40 cycles of 95°C 10s and 60°C 30s. The absolute quantification of the N1 copy number was interpolated using a standard curve with 10^7^-10^1^ serial 10-fold dilution of a control plasmid (nCoV-CDC-Control Plasmid, Eurofins).

### VeroE6 Plaque Assay

Virus stocks with a pre-determined virus copy number were evaluated in a conventional VeroE6 plaque assay to determine if the virus titers obtained using both methods correlate. 4×10^5^ VeroE6 cells (ATCC) were plated in 6-well plates and allowed to adhere overnight at 37°C. Cells were washed once with PBS and infected with 1 ml of viral stocks serially diluted in 2× MEM complete media (10% FBS, 20 mM HEPES, 2× Pen-Step, 2× NEAA and 2× Sodium Pyruvate) for 1 hr at 37°C. After infection, 1 ml of sterile 2.5% cellulose overlay solution (Sigma, Cat. No. 435244-250G) was added to each well and mixed thoroughly. Cells were incubated at 37°C for an additional 48 hr before the media/overlay was removed and the cells fixed in 4% paraformaldehyde (PFA) for 10 min at room temperature. The PFA was removed and the cells were stained with 1% crystal violet in ethanol for 1 minute and washed three times with distilled water. Plaques were manually counted from each well.

### Immunofluorescence Assay

Primary human airway epithelial cells fully differentiated in air-liquid interface cultures [59] were infected with different SARS-CoV-2 variants with or without IFNβ. The apical surface was washed with culture medium daily for quantitative PCR. At 96 h post-infection, the cultures were fixed with 4% PFA and wholemount labeled with anti-Spike antibody (Clone ID007, Cat. No. 40150-R007, Sino Biological) followed by Alexa-Dye conjugated secondary antibody. An LSM 900 confocal microscope (Zeiss) was used to generate composite images of the entire culture surface. Spike+ cells were enumerated using the Cell Counter plugin in the ImageJ Software (NIH).

### Antiviral inhibition assay

We used a non-saturating dose of the amplified virus stock for the IFN inhibition assays. These titers were expected to yield ~10^5^ copies per 5 μl input RNA extract (S1B Fig). Recombinant IFNs were obtained from PBL Assay Science. These recombinant IFNs were assayed to be >95% pure by SDS-PAGE according to the manufacturer. In addition to the IFN-Is (12 IFNα subtypes, IFNβ and IFNω), we also evaluated 3 IFNλ subtypes (IFNλ1, IFNλ2, IFNλ3). To normalize the IFNs, we used molar concentrations [23] instead of international units (IU), as IU values were derived from inhibition of encelphalomyocarditis virus, which may not be relevant to SARS-CoV-2. Importantly, molar concentrations were used to normalize the relative signaling potencies of the IFNα subtypes and IFNβ [23, 24]. To find a suitable dose to screen 17 IFNs in parallel, we performed a dose-titration experiment of the USA-WA1/2020 strain with IFNβ and IFNλ1. A dose of 2 pM allowed for maximum discrimination of the antiviral potency IFNβ versus IFNλ1 (S1C Fig). Thus, this dose should be within the dynamic range of inhibition of the diverse IFNs investigated. Serial 10-fold dilutions of IFNβ and IFNλ1 were also used in follow-up experiments. Thus, in 48-well plates, we pre-incubated 2.5×10^4^ A549-ACE2 cells with the IFNs for 18 h, then infected with the A549-amplified virus stock for 2 h. After two washes with PBS, 500 μl complete media containing the corresponding IFNs were added. The cultures were incubated for another 24 h, after which, supernatants were harvested for RNA extraction and qPCR analysis. A similar procedure was employed for Calu-3 cells, except that IFNλ1 was replenished at 2 dpi and supernatants harvested at day 3.

### Statistical analyses

Data were analyzed using GraphPad Prism 8. Differences between the IFNs were tested using a nonparametric two-way analysis of variance (ANOVA) followed by a multiple comparison using the Friedman test. Pearson correlation coefficients (R^2^) values were computed for linear regression analyses. Paired analysis of two isolates against multiple IFNs were performed using a nonparametric, two-tailed Wilcoxon matched-pairs rank test. Differences with *p*<0.05 were considered significant. Nonlinear regression curves were used to fit using either a one-site total or two-phase exponential decay equation on log-transformed data.

## Acknowledgments

We thank Cara Wilson, Ulf Dittmer and Kathrin Gibbert for scientific advice; Mercedes Rincon and Elan Eisenmesser for assistance with construction and characterization of the A549-ACE2 cells; Zach Wilson, Jill Garvey, Stephanie Torres-Nemeti, Brett Haltiwanger and Marcia Finucane for Biosafety Level-3 infrastructure support; and Roman Wölfel, Rosina Ehmann, Adolfo García-Sastre, Alex Sigal, Tulio de Oliveira, Bassam Hallis, Matsuyo Takayama-Ito, Richard Webby, Anami Patel, Cathleen Seager, BEI Resources (NIAID) and the CDC for the SARS-CoV-2 isolates.

