## Supplemental Table 1 and Figures 1-6 for "Interferon Resistance of Emerging SARS-CoV-2 Variants"

**This PDF file includes:**

Supporting Data Table 1
Supporting Data Figures 1 to 6

**Supporting Table 1. SARS-CoV-2 Isolates Tested for IFN-I and IFN-III sensitivity.**

| BEI Catalogue Number | Isolate Name | Collection Date | GISAID clade | Lineage (WHO) | GISAID accession number | Source | Amino acid Mutations* | Depositor | BEI Resources Lot Number | Notes |
|---|---|---|---|---|---|---|---|---|---|---|
| NR-52281 | USA-WA1/2020 | Jan 19, 2020 | S | A | EPI_ISL_404895 | Male in 30s with mild disease, returning traveler from Wuhan, China | NS8 L84S | Centers for Disease Control and Prevention, Atlanta, GA, USA | 70036318 | Standard strain used in multiple in vitro and in vivo SARS-CoV-2 studies |
| NR-52370 | Germany/BavPat1/2020 | Jan 28, 2020 | G | B | EPI_ISL_406862 | Mildly asymptomatic adult male from a Bavarian transmission cluster | Spike **D614G** | Drs. Roman Wölfel and Rosina Ehmann, Bundeswehr Institute of Microbiology, Munich, Germany | 70036595 | Also referred to as Germany/MUC-IMB1/2020 or Human/DEU/BavPat-ChVir929/2020. Spike mutation was associated with increased transmissibility. |
| NR-53514 | New York-PV08410/2020 | Mar 16, 2020 | GH | B.1 | EPI_ISL_421374 | Isolated from a nasal swab collected from a 63 year old male patient with a fatal respiratory illness in New York, USA | NSP2 T85I, NSP12 P323L, Spike **D614G**, NS3 Q57H | Dr. Adolfo García-Sastre, The Icahn School of Medicine at Mount Sinai Medical School, New York, NY, USA | 70036345 | Belongs to clade 'A2a', a large transmission cluster in New York City during a major outbreak in March 2020. It was phylogenetically similar to isolates from Europe. |
| NR-54008 | hCoV-19/South Africa/KRISP-EC-K005321/2020 | Nov 15, 2020 | GH | B.1.351 (Beta) | EPI_ISL_678570 | Isolated from an oropharyngeal swab from a 57-year-old human male in Harry Gwala district, KwaZulu-Natal, South Africa. | Spike A243del, A701V, D80A, D215G, **D614G**, **E484K**, **K417N**, L242del, L244del, **N501Y**; E P71L; N T205I; NS3 Q57H, S171L; NSP2 T85I; NSP3 K837N; NSP5 K90R; NSP6 F108del, G107del, S106del; NSP12 P323L | Dr. Alex Sigal, Africa Health Research Institute and Prof. Tulio de Oliveira, KwaZulu-Natal Research Innovation and Sequencing Platform (KRISP), Durban, South Africa | 70041939 | The deposited virus (after passage 3) harbored additional mutations compared to the clinical isolate: deletion in Furin cleavage site in Spike (677-681del), ORF1a (Q3878R) |
| NR-54011 | USA/CA_CDC_5574/2020 | Dec 29, 2020 | GRY | B.1.1.7 (Alpha) | EPI_ISL_751801 | Isolated from nasopharyngeal swab in San Diego, California, USA. Lineage first reported in the United Kingdom. | Spike A570D, **D614G**, D1118H, H69del, **N501Y**, P681H, S982A, T716I, V70del, Y145del; M V70L; N **D3L**, G204R, R203K, S235F, NS3 T223I; NS8 Q27stop, R52I, Y73C, A890D; NSP3 I1412T, T183I; NSP6 F108del, G107del, S106del; NSP12 P323L, A454V, K460R | Centers for Disease Control and Prevention, Atlanta, GA, USA | 70041598 | One additional SNP in ORF1ab L3826F was reported in the deposited passage two virus, in comparison to the clinical specimen. |
| NR-54008 | hCOV-19/ England/204820464/2020 | Nov 24, 2020 | GRY | B.1.1.7 (Alpha) | EPI_ISL_683466 | Isolated from a 58-year old male from England, United Kingdom | Spike A570D, **D614G**, D1118H, H69del, **N501Y**, P681H, S982A, T716I, V70del, Y145del; N **D3L**, G204R, R203K, S235F, NS8 R52I, Y73C; NSP3 A1305V, I1412T, T183I; NSP6 F108del, G107del, S106del; NSP12 P323L; NPS13 K460R; NSP14 E347G | Dr. Bassam Hallis, Pre-Clinical Development at Public Health England, Salisbury, UK | 70041933 | Also referred to as UK/VUI/3/2020 isolate. |
| NR-54892 | hCoV-19/Japan/TY7-503/2021 | Jan 6, 2021 | GH | P.1 (Gamma) | EPI_ISL_792683 | Isolated in airport quarantine in Japan from a COVID-19 positive passenger from Brazil | Spike D138Y, **D614G**, **E484K**, H655Y, **K417T**, L18F, **N501Y**, P26S, R190S, T20N, T1027I, V1176F; N G204R, P80R, R203K; NSP3 S253P, K977Q, S370L; NSP8 E92K; NSP3, NSP3; NSP6 F108del, G107del, S106del; NSP12 P323L; NSP13 E341D | Dr. Matsuyo Takayama-Ito, National Institute of Infectious Diseases, Japan | 70042875 | The sequence of this isolate branched off from the B.1.1.28 lineage. The deposited virus (Passage 2 in Vero E6/TMPRSS2 cells) was reported to have an additional NSP6 F184V mutation as compared to the clinical isolate. |
| NR-55611 | hCoV-19/USA/PHC658/2021 | May 2021 | GH | B.1.617.2 (Delta) | | Isolated from a patient with severe COVID-19 in Memphis, TN | Spike T19R, K77T, FR157-158del, **L452R**, T478K, **D614G**, P681R, D950N; NSP2 D84N, P129L; NSP3 P822L; NSP4 D217N, F375S; NSP6 H11Q; NSP12 P323L; NSP15 K259R; NS3 S26L; M I82T; N D63G, R203M, D377Y; ORF7a 44-100fs, T120I; ORF8 C102F | Drs. Richard Webby and Anami Patel, St. Jude's Hospital, Memphis, TN, USA | 70045238 | Variant was first detected in India, and accounts for majority of infections in the USA as of July 2021. **ORF7a was deleted.** Additional mutations in Spike G142D and N P142Q were detected in the deposited virus compared to the clinical isolate. |

*Mutations were inferred relative to the reference hCOV-19/Wuhan/WIV04/2019 sequence (GISAID.org).

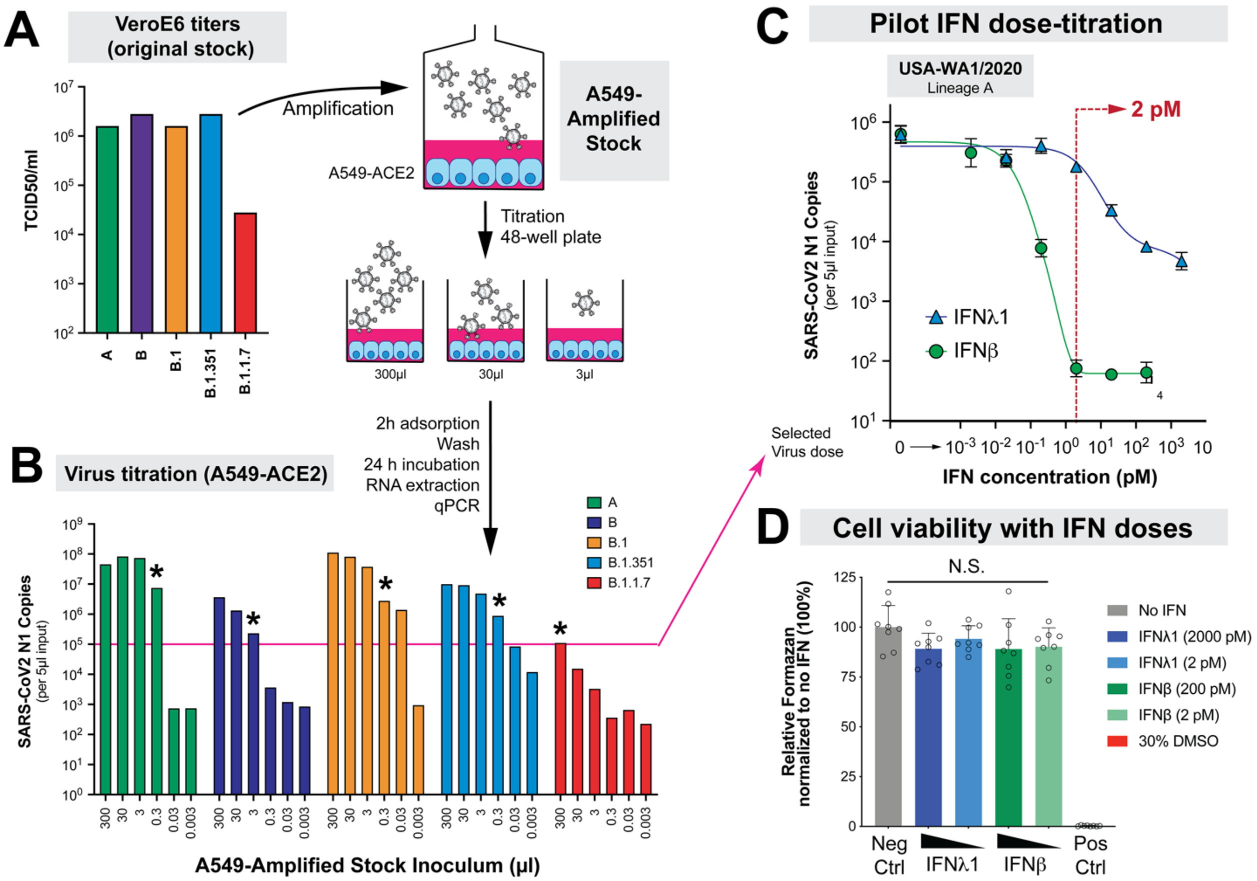


**S1 Figure. Preparation of SARS-CoV-2 Isolates for IFN sensitivity assays.** (A) SARS-CoV-2 stocks were obtained from BEI Resources. These original stocks were propagated in VeroE6 cells and titered to obtain 50% tissue culture infectious doses ($TCID_{50}$) per ml. Upon receipt, these original stocks were passaged and amplified once in A549-ACE2 cells. (B) Amplified virus stocks were titered in A549-ACE2 cells in a 48-well format for the subsequent inhibition assays. We aimed to infect cells to yield ~$10^5$ copies, which would provide a 3-log dynamic range of inhibition. The pink line corresponds to this target virus copy number in untreated cells (~$10^5$ copies) and asterisks correspond to the approximate amount of virus that was used to reach that level for the inhibition assays. Not shown in the graphs is the P.1 isolate which had an initial titer of $2.8 \times 10^6$ $TCID_{50}$/ml and upon low-passage amplification reached >$10^5$ copies in A549-ACE2 cells. (C) Identification of an IFN dose for quantitative assessments of variant IFN-sensitivity. A ten-fold dilution series of IFNλ1 (0.02 to 2000 pM) and IFNβ (0.002 to 200 pM) were pre-incubated with A549-ACE2 cells for 24 h and USA-WA1/2020 RNA copies were evaluated by qPCR. Red dotted lines correspond to the dose (2 pM) that allowed for maximum discrimination of the antiviral potencies of the 2 IFNs. Points correspond to mean values and error bars correspond to standard deviation of triplicate measurements. (D) The viability of A549-ACE2 cells was assessed 36 h post-treatment with the maximum and 2 pM dose of IFNλ1 and IFNβ using the MTT assay. Error bars correspond to SD of 8 replicates. Data were normalized to the negative (media only) control. 30% DMSO was used as a positive control for cell death. Data were analyzed using a one-way ANOVA and a post-hoc Tukey's multiple comparison test. All pairwise comparisons between the bars under the line, 'NS' were not significant ($p>0.05$).

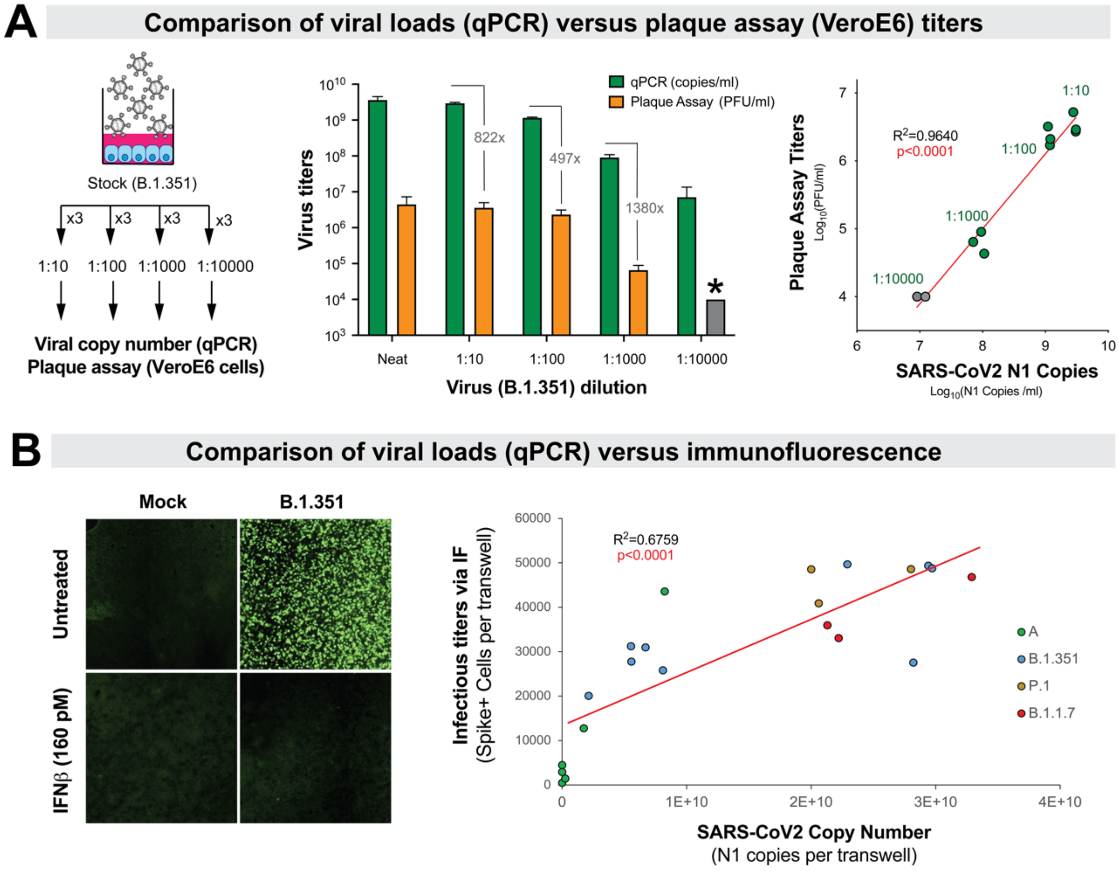


**S2 Figure.** SARS-CoV-2 N1 copy numbers significantly correlate with infectious titers. (A) Comparison of virus titers obtained using qPCR versus the VeroE6 plaque assay. (*Left*) Four 10-fold dilutions of a virus stock (B.1.351) were prepared in triplicate and subjected to both virological assays. (*Middle*) Mean values ± SD for these titrations are depicted in the bar graphs. *Infectious titers were below $10^4$ PFU/ml. As the values were saturated at the highest virus concentration, (*Right*) linear regression excluded the neat dilution. One outlier qPCR value (log copy number=4.15 versus 7.98 and 8.02) at 1:10,000 was removed. Note that a significant correlation was still observed even if the 1:10,000 dilution was removed ($R^2$=0.94, p<0.0001). Based on the ratio of copy number versus plaque forming units (*Middle*), we estimate that 1 plaque forming unit corresponds to ~900 SARS-CoV-2 N1 copies. (B) Primary human airway epithelial cell cultures air-liquid interface cultures (grown on Transwell culture inserts) were infected with different SARS-CoV-2 variants with or without IFNβ. At 96 h, supernatants were harvested for RNA extraction and qPCR, and the cells were fixed then labeled with an anti-Spike antibody for immunofluorescence microscopy. (*Left*) A representative en face view of the Transwell culture surface is shown. Green foci correspond to Spike+ cells. (*Right*) Spike+ cell numbers were compared to supernatant virus copy number from the same Transwell cultures by linear regression.

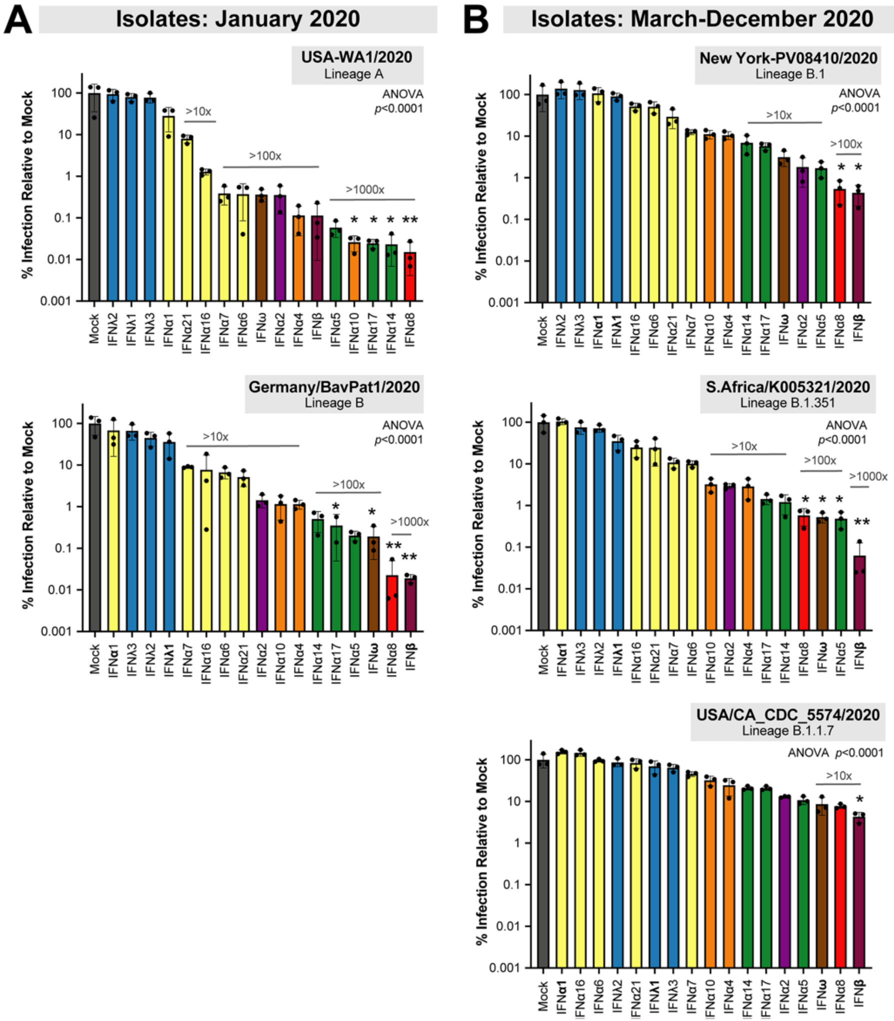


**S3 Figure. Sensitivity of SARS-CoV-2 strains to IFN-I and IFN-III interferons.** Viral RNA load data in Fig. 2 were normalized to mock (no IFN) as 100% and plotted in log-scale. Each dot corresponds to one of triplicate experiments, bars correspond to mean values and error bars correspond to standard deviations. The IFN sensitivity profiles were subdivided between isolates collected (A) early; and (B) later in the COVID-19 pandemic. Differences between the IFNs were evaluated using Friedman test. The *p*-values were noted in each graph. A post-hoc multiple comparisons test was performed for each IFN against the mock control; *, $p<0.05$, **, $p<0.01$.

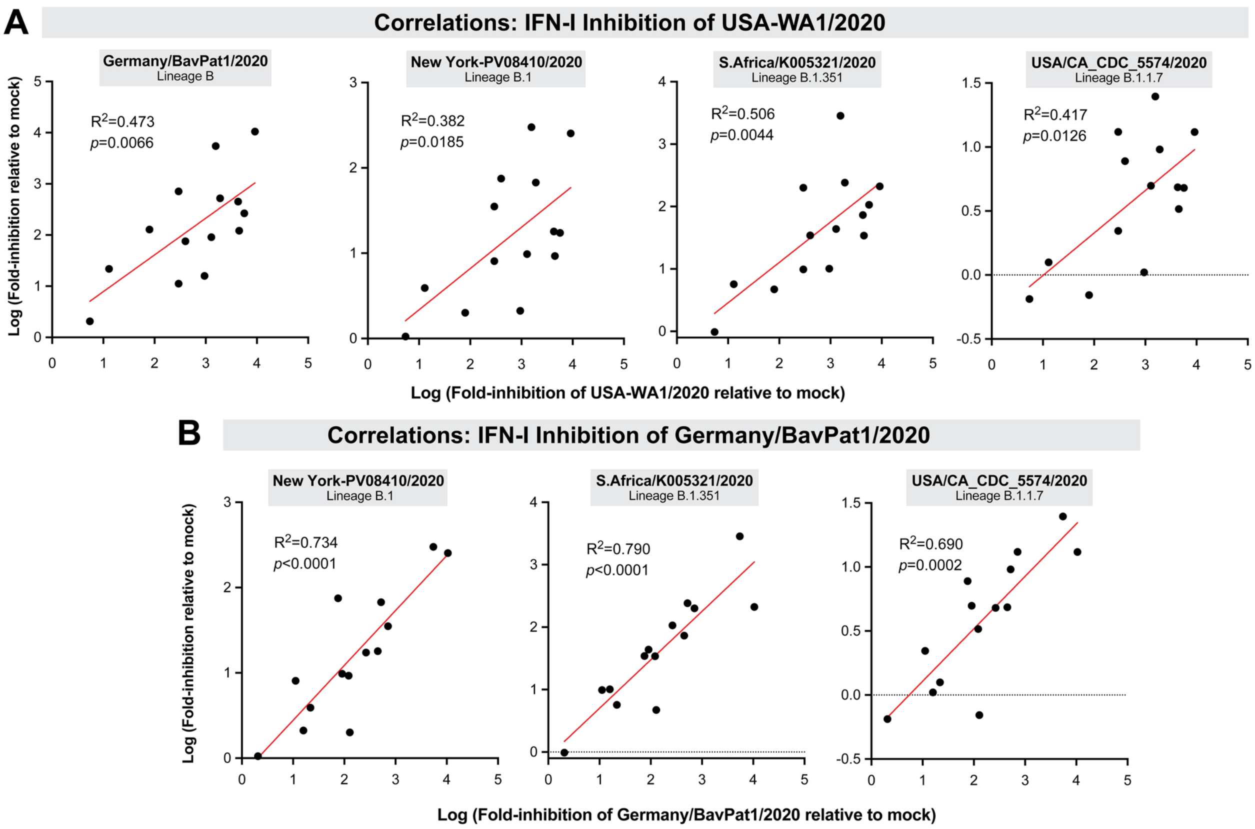


**S4 Figure. Correlation between IFN-I inhibition of different SARS-CoV-2 isolates.** Fold-inhibition values relative to mock were calculated for each IFN-I tested then compared to (A) the USA-WA1/2020 strain and (B) the Germany/BavPat1/2020 strain. Log-transformed values were compared. Linear regression was performed in GraphPad Prism 8, with the best-fit line (red), $R^2$ and *p*-values indicated.

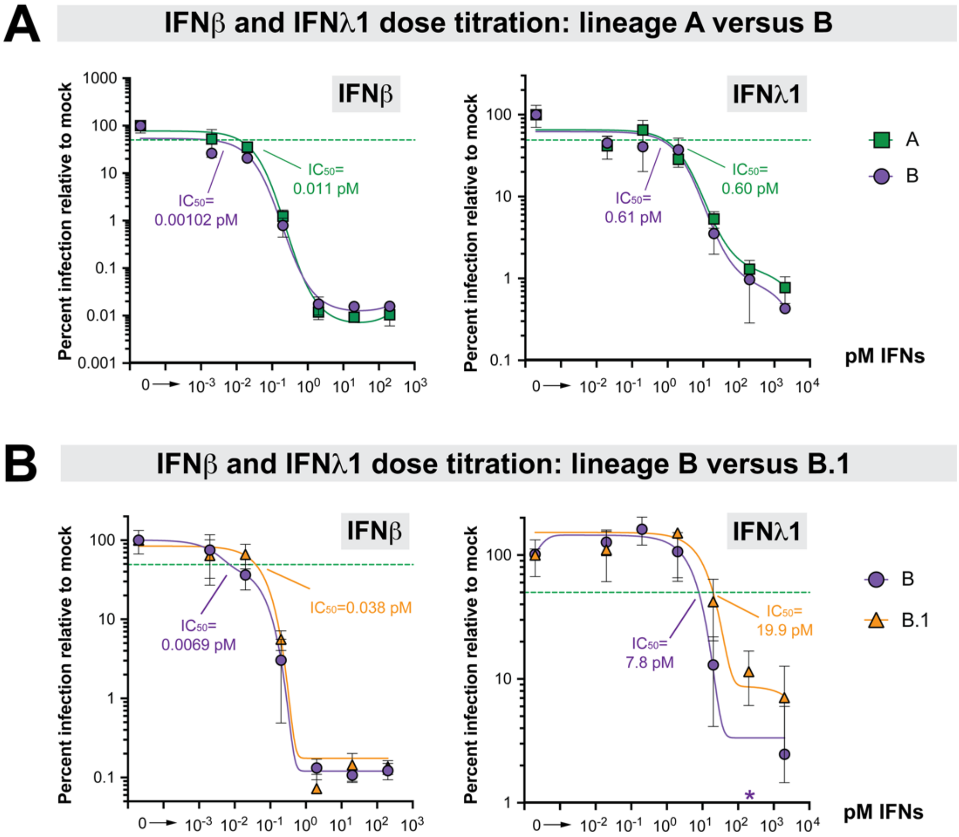


**S5 Figure. IFN-I resistance of early SARS-CoV-2 isolates.** Dose-titration of IFNβ and IFNλ1 against lineage B (Germany/BavPat1/2020) versus (A) lineage A (USA-WA1/2020) and (B) lineage B.1 isolate (New York-PV08410/2020). A549-ACE2 cells were pre-treated with serial 10-fold dilutions of IFNs for 18 h in triplicate and then infected with SARS-CoV-2. Supernatants were collected after 24 h, SARS-CoV-2 N1 gene copy numbers were determined by qPCR in triplicate, and then the mean copy numbers were normalized against mock as 100%. Non-linear best-fit regression curves of mean normalized infection levels were used to interpolate 50% inhibitory concentrations (green dotted lines). *Note that the value for lineage B at 200 pM IFNλ1 was 0.54, precluding efforts to find a best-fit curve for IC50 determination; this datapoint was therefore not included in the curve-fitting.

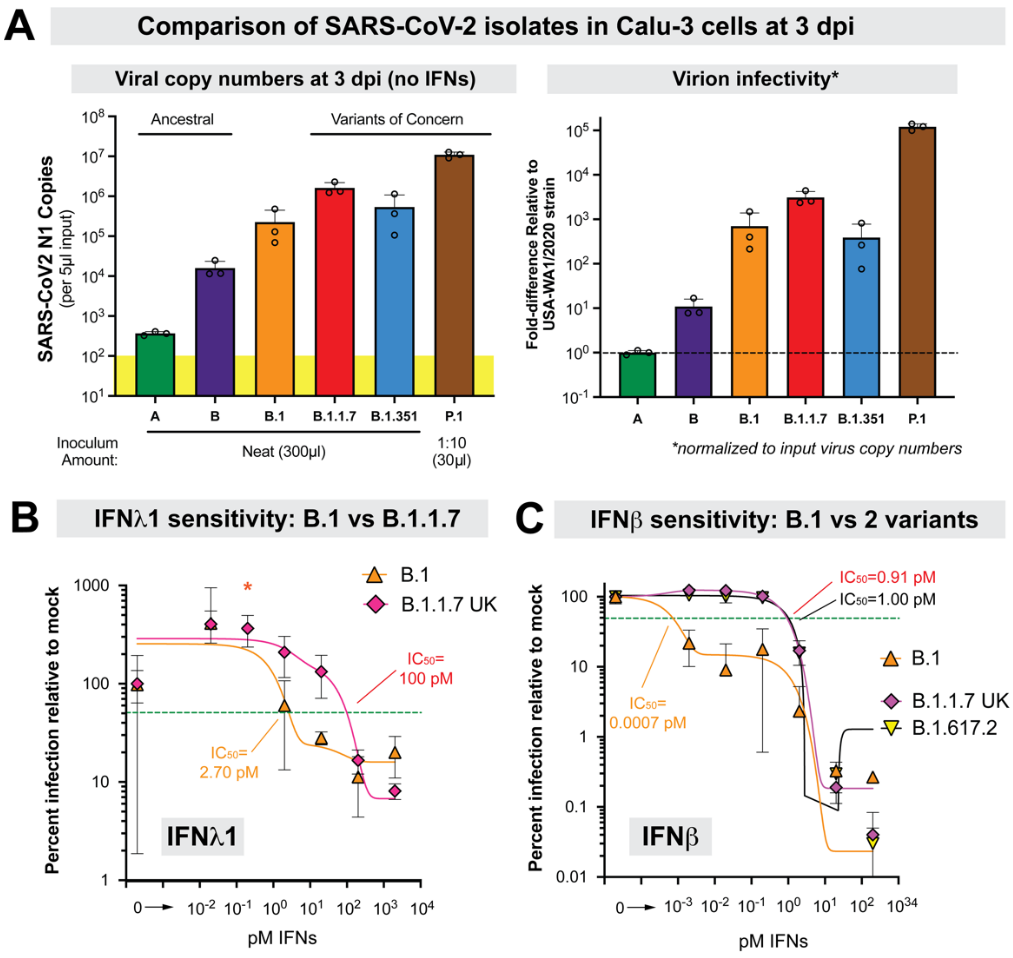


**S6 Figure. Evaluation of SARS-CoV-2 isolates in Calu-3 cells.** (A) Susceptibility of Calu-3 cells to infection with diverse SARS-CoV-2 isolates. Please refer to S1 Table for the specific strains used for each lineage. B.1.1.7 corresponds to the England/204820464/2020 isolate. (*Left*) SARS-CoV-2 N1 gene copy numbers were quantified at 3 dpi. The yellow bar corresponds to the assay limit of detection. (*Right*) The infectious titers at 3 dpi from the left panel was divided by the N1 copy number of the virus inoculum, then normalized against the lineage A isolate (USA-WA1/2020). This ratio provides an estimate of the infectivity of the isolates. Ancestral isolates replicated poorly in Calu-3 cells, and thus were not optimal for IFN treatment studies due to the very low dynamic range. Comparable titers were observed for B.1, B.1.1.7 and B.1.351, whereas the P.1 isolate was highly infectious in this cell line. (B) Comparison of IFNλ1 sensitivity of B.1 versus B.1.1.7. Calu-3 cells were treated with varying doses of IFNλ1 at -1 and 2 dpi. (C) Comparison of IFNβ sensitivity of B.1 versus B.1.1.7 and B.1.617.2. Calu-3 cells were treated with varying doses of IFNβ at -1 dpi. For both (B) and (C), supernatants were collected at 3 dpi for qPCR evaluation. Error bars correspond to the mean of 3 biological replicates; each biological replicate was evaluated by qPCR in triplicate. Nonlinear regression curves to calculate IC50s were generated using GraphPad Prism 9. *The titer for B.1 at 0.2 pM showed 9.2-fold higher infection than mock, precluding efforts to find a best-fit curve for IC50 determination. This datapoint was therefore not included in the curve-fitting.
